# The Biological Basis of Mathematical Beauty

**DOI:** 10.1101/367185

**Authors:** Semir Zeki, Oliver Y. Chén, John Paul Romaya

**Affiliations:** Laboratory of Neurobiology, Division of Cell & Developmental Biology, University College London, London WC1E 6BT; Laboratory of Neurobiology, University College London, London, UK; Department of Psychology, Yale University, New Haven, CT, USA

**Author notes:** Corresponding author: Semir Zeki.

## Abstract

Through our past studies of the neurobiology of beauty, we have come to divide aesthetic experiences into two broad categories: biological and artifactual. The aesthetic experience of biological beauty is dictated by inherited brain concepts, which are resistant to change even in spite of extensive experience. The experience of artifactual beauty on the other hand is determined by post-natally acquired concepts, which are modifiable throughout life by exposure to different experiences (Zeki, 2009). Hence, in terms of aesthetic rating, biological beauty (in which we include the experience of beautiful faces or human bodies) is characterized by less variability between individuals belonging to different ethnic origins and cultural backgrounds or the same individual at different times. Artifactual beauty (in which we include the aesthetic experience of human artifacts such as buildings and cars) is characterized by greater variability between individuals belonging to different ethnic and cultural groupings and by the same individual at different times. In this paper, we present results to show that the experience of mathematical beauty (Zeki *et al* 2014), even though it constitutes an extreme example of beauty that is dependent upon (mathematical) culture and learning, belongs to the biological category and obeys one of its characteristics, namely a lesser variability in terms of the aesthetic ratings given to mathematical formulae experienced as beautiful.

## Introduction

In an earlier study (Zeki *et al*. 2014), we reported that the experience of mathematical beauty (by mathematicians) correlated with activity in field A1 of the medial orbito-frontal cortex (mOFC); moreover, the intensity of activity there was parametrically related to the declared intensity of the beauty experienced by the mathematicians when viewing the mathematical formulae. This was a somewhat surprising result, at least to us. The experience of mathematical beauty is derived from a highly intellectual, cognitive source; it is perhaps the most extreme example of aesthetic experience that is dependent upon culture and learning. Unlike visual or musical beauty, only those versed in mathematics can experience the beauty of mathematical formulations. And yet the experience of mathematical beauty correlates with activity in the same part of the emotional brain as the experience of beauty derived from sensory sources, such as the visual or the musical (Ishizu and Zeki 2011). This naturally leads one to enquire further into the nature of mathematical beauty and whether it can be classified as belonging to the category of biological beauty.

In pursuing our studies of beauty, we thought it interesting to add to our previous study of the experience of mathematical beauty by analysing our results further, with the following questions in mind: what was the degree of variability in the ratings given to equations that had been rated as beautiful; did that variability differ in any significant way from the variability in the ‘non-beautiful’ ratings that had been assigned to other equations. Our only hypothesis in this regard was that, if mathematical beauty belongs to the biological category, then there should be significantly less variability among equations given high ratings than among others. We indeed found this to be the case, which reinforced our view that mathematical beauty belongs to the category of biological beauty, for reasons which we explore more fully in the Discussion section.

## Material and methods

A full description of the subjects and methods used to rate mathematical equations is given in Zeki *et al*. (2014), where all the 60 mathematical formulae used in the study are also tabulated. In brief, 15 mathematicians (3 females, in the age range of 22–32 years) and all of them postgraduate students or post-doctoral fellows in mathematics took part in the study. Each was given 60 mathematical equations to study at leisure and rate according to the aesthetic experience aroused in them on a scale of −5 (ugly) to +5 (beautiful). Subsequent to a brain scanning experiment, to determine the brain areas in which activity correlates with the experience of mathematical beauty (the results of which are reported in Zeki *et al*., 2014), each subject was asked to report their level of understanding of each equation on a scale of 0 (no understanding) to 3 (profound understanding) and to report their emotional reactions to the equations. In this paper, we deal only with the ratings given and do not consider the results of the brain imaging study. The ratings used appear in Data Sheet 3 of the supplemental data in tables 1 (pre-scan beauty ratings) and 6 (post-scan understanding ratings) of Zeki *et al*. (2014).

## Results

We undertook the following statistical analyses of the pre- and post-scan ratings given to the mathematical formulae by in Zeki *et al* (2014), using the notations given below:

*r_ij_* denotes the **beauty rating** that the *i^th^* subject gives to the *j^th^* formula.

*u_ij_* denotes individual *i’s* **understanding** of the *j^th^* formula.

*N* denotes the total number of subjects.

*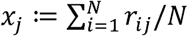* and 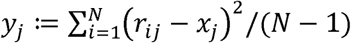 represent the mean beauty-rating **[m-BR]** and the standard deviation of beauty ratings **[sd-BR]**, respectively, given to the *j^th^* formula across subjects.

*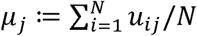* and 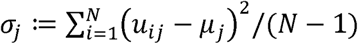 represent the mean understanding rating of the formula **[m-UR]** and standard deviation of the understanding rating **[sd-UR]**, respectively, given to the *j^th^* formula across subjects.

1. We calculated the m-BR and the sd-BR for each formula across subjects. This yields 60 m-BR and 60 corresponding sd-BR values **[see Figure 1]**. The graph has a pronounced negative trend, showing that there was generally a lower standard deviation for formulae rated as beautiful compared to ones rated as ugly [Pearson *r*: −0.59, *p* < 10^−6^; Spearman *ρ*: −0.53, *p* < 10^−4^). In simpler terms, there was a higher consensus among our sample of 15 subjects about the beautiful equations than about the not beautiful ones since, unlike the equations rated as beautiful, there was greater variability for those rated as not beautiful.
2. We sought to uncover the linear relationship between the m-BR and the sd-BR across equations. Formally, consider 

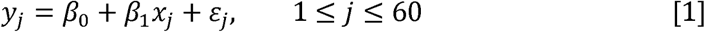

where *x_j_* and *y_j_*- refer to the m-BR and the sd-BR of the *j^th^* equation, *β*_0_ and *β*_1_ are parameters for the intercept and slope, and *ε_i_* indicates the residual term (*i.e*., the information not explained). Our data shows that the estimate for *β*_1_ or 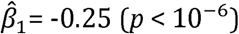. In other words, if a formula is on average rated one point higher than a second formula, the standard deviation of the ratings across subjects (which quantifies the disagreement among subjects) for the former formula is 0.25 units less than the standard deviation of the ratings for the latter formula.
3. Although a small sd-BR is significantly associated with a large m-BR (*i.e*. the more beautifully rated formulae have smaller standard deviations), it remained possible that these are confounded by subjects’ understanding of the mathematical formulae. To check for confounds, we reran Model (1), where the regressor *x_j_* (the m-BR of formula *j*) was replaced with *z_j_* (the m-UR of formula *j*) **(see Figure 2 (a))**. Our data showed that the m-UR does not significantly affect the sd-BR (Pearson *r*: −0.25, *p* >0.05; Spearman *ρ*: −0.24, *p* >0.05). Similarly, we show that that the sd-UR does not significantly affect the sd-BR (Pearson *r*: 0.30, *p* >0.05; Spearman *ρ*: 0.29, *p* >0.05) **(see Figure 2 [b])**.
4. This far, we have shown that the m-UR did not significantly affect the s-BR; the possibility remains that the m-UR and the m-BR may be jointly related to the s-BR. We therefore ran tests to learn if adding the m-UR would improve the prediction of the s-BR. Formally, we compared Model (1) with the following model 

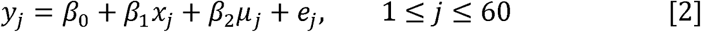

where *μ_j_* denotes the m-UR of the *j^th^* formula, and *e_j_* indicates the residual term. An *F*-test statistic is defined as 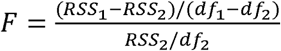, where *RSS*_1_ and *RSS*_2_ denote residual sums of squares for Models (1) and (2), respectively, and *df*_1_ and *df_2_* denote degrees of freedom (*i.e*. number of data points minus number of parameters) for Models (1) and (2), respectively (Fisher, 1925)(Ott, R. Lyman, 1980)(Bickel & Doksum, 2000). The *F*-test can be used for determining whether a "more complicated” model with additional parameters (*e.g*. Model [2]) is *significantly* better than a baseline model (*e.g*. Model [1]). Using an *F*-test, we show that adding variable (*μ*) in Model [2] does not significantly reduce prediction errors in Model [1] (F=0.04, p=0.84). The *F*-test result suggests that including the m-UR (*μ*) does not add additional information to the established association between the sd-BR (*y*) and the m-BR (*x*) in Model [1]. Similarly we can construct a further model

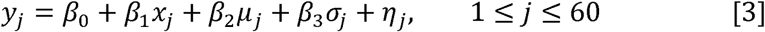

where *σ_j_* denotes the sd-UR of the *j^th^* formula, and *η_j_* indicates the residual term. Comparing Models (1) and (3) we show that including both the m-UR (*μ*) and the sd-UR (*σ*) of the formulae does not add additional information into the established association between the sd-BR (*y*) and the m-BR (*x*) (F=1.59, p=0.21).
5. Additionally, we show that including two-way interactions (m-BR × m-UR, m-BR × sd-UR, and m-UR× sd-UR) and three-way interactions (m-BR × m-UR × sd-UR) (Fisher, 1936) (Wu & Hamada, 2000) does not add additional information to the established association between the standard deviation of the beauty rating (*y*) and the mean beauty rating (*x*) in Model (1) (*F*=0.49, *p*=0.75).
6. Taken together, our analysis shows that high mean rating of formulas in a population is significantly (negatively) associated with the standard deviation of the ratings. Specifically, one unit increase of mean rating leads to −0.25 units decrease of standard deviation of a formula. Further, such association is neither due to, nor can be further explained by, one’s understanding of the formulas, suggesting that there is a unified aesthetical appreciation of mathematics among individuals, and that such aesthetical appreciation is not necessarily related to one’s understanding of mathematics.

**Figure 1.**
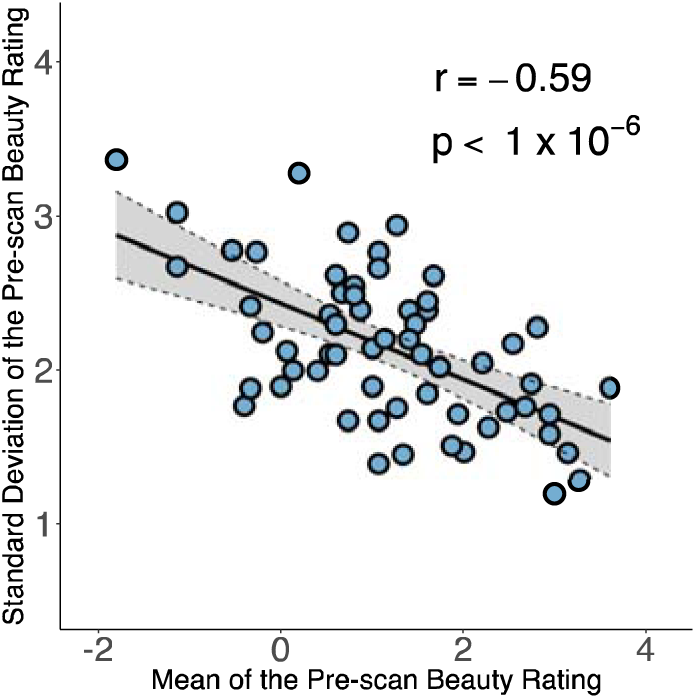
A plot of the mean beauty rating [m-BR) for each equation against the standard rating [sd-BR) for the ratings given to each equation. Pearson *r*: −0.59, *p <* 10^−6^; Spearman *ρ*: −0.53, *p <* 10^−4^). Each circle corresponds to one equation; its value on the horizontal line indicates the average beauty ratings of the equation across all subjects and its value on the vertical line indicates the standard deviation of the beauty ratings. Grey area indicates the 95% confidence band for best-fit line.

**Figure 2:**
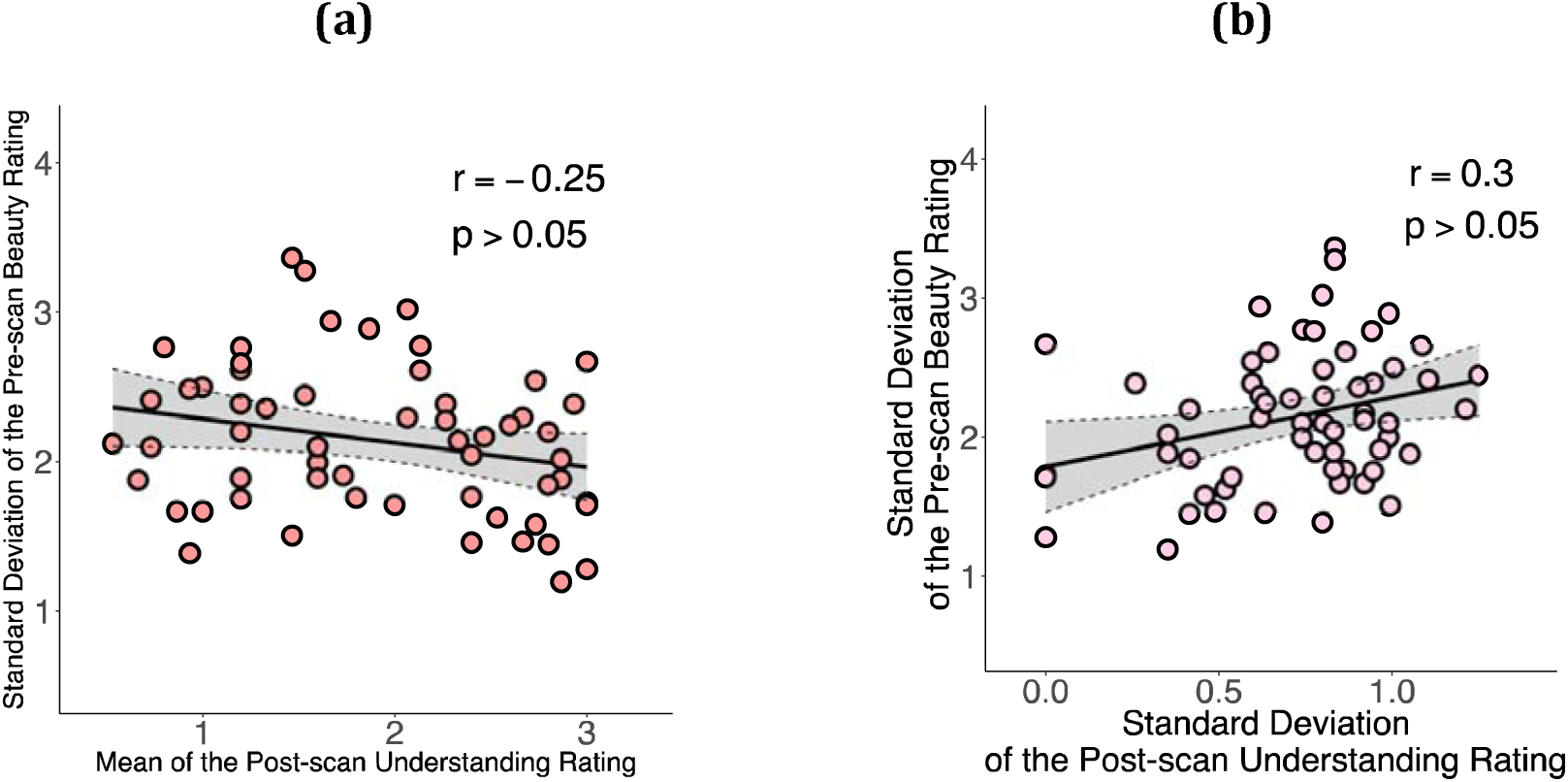
(a) A graph relating the standard deviation of beauty rating (sd-BR) for each equation against mean understanding rating (m-UR) for each equation (Pearson *r*: −0.25, *p* >0.05; Spearman *ρ*: −0.24, *p* >0.05). Each circle corresponds to one equation; its value on the horizontal line indicates the average understanding ratings of the equation across all subjects and its value on the vertical line indicates the standard deviation of the beauty ratings. Grey area indicates the 95% confidence band for best-fit line. (b) A graph relating the standard deviation of the beauty rating (sd-BR) to that of the understanding rating (sd-UR). (Pearson *r*: 0.30, *p* >0.05; Spearman *ρ*: 0.29, *p* >0.05). Here, each circle corresponds to one equation; its value on the horizontal line indicates the standard deviation of the understanding ratings of the equation across all subjects and its value on the vertical line indicates the standard deviation of the beauty ratings. Grey area indicates the 95% confidence band for best-fit line.

## Discussion

That the experience of mathematical beauty correlates with activity in the same part of the emotional brain as the experience of beauty derived from other sources, including sensory and moral ones(Ishizu & Zeki, 2011), raises the question of how to classify mathematical beauty.

We have in the past suggested that the classification of experiences, ranging from ordinary sensory ones such as that of colour, to aesthetic experiences such as that of beauty, can be subdivided into two broad categories. At one end are experiences belonging to the *biological* category: these have a biologically inherited brain concept as a basis (Zeki 2009). Such biological concepts lead to experiences that are shared universally, by all ethnic groups and are independent of learning. Moreover, they are not easily modifiable by experiment or repeated exposure to a variant that departs significantly from the inherited concept, at least within the limits tested (see Chen and Zeki 2011). This entitles a subject having an aesthetic experience, which belongs in the biological category and determined by an inherited brain concept to suppose that the experience is similar to the one that others would have in similar circumstances and that it has, therefore, universal validity and assent (see also Zeki and Chen 2017). A prime example of this is in the experience of colour categories. The colours of objects and surfaces remain constant in spite of wide fluctuations in the wavelength-energy composition of the light reflected off them (Land 1974). This phenomenon is generally referred to as colour constancy although we much prefer the term constant colour categories (see Zeki *et al*. 2017). This is because, although the colour category does not change with such fluctuations, the exact hue (shade of colour) of a patch belonging to a given colour category will do so. The generation of constant colour categories is due, we suppose, to an inherited brain program, which compares the wavelength-energy composition of light coming from one patch with that coming from surrounding patches, thus generating ratios between the two for every waveband (Land, 1974), with the ratios remaining constant in spite of significant changes in the amounts of light of different wavebands reflected from the viewed patch and its surrounds. In fact, psychophysical experiments that we have undertaken (in preparation) demonstrate that there is very little variability among humans of different ethnic and cultural backgrounds when asked to assign the colour of patches (when the wavelength-energy composition of the light reflected from them changes) to a standard set of coloured chips. This is because, due to an inherited brain program for generating constant colour categories, the patches do not change their colour categories with such fluctuations. Hence humans can (and do) suppose that others have a similar colour experience to them and that their experience has, therefore, universal, assent.^3^

At the other end are experiences determined by acquired brain concepts, examples being that of man-made artefacts such as buildings or cars or vases and a variety of manufactured goods. The concept underlying these experiences are acquired post-natally and are modifiable throughout post-natal life (Zeki 2009), which can be demonstrated experimentally (Chen and Zeki 2011). Because they are based on individual experience and experimentation, the experiencing subject cannot assume that others will have the same experience. As an example, someone who is brought up in a particular architectural environment, say a Western one, cannot assume that another human from a different cultural environment will find the same satisfaction in Western architecture. Moreover, since the brain concept itself is acquired post-natally and changes with new experiences, one cannot even assume that an aesthetic judgment on the architectural merit of a building made today will be the same as the one made in the past or that will be made in the future (Zeki and Chén 2016).

### The categorization of mathematical beauty

This naturally raises the question of whether the experience of mathematical beauty belongs in the biological or the artifactual category.

The experience of mathematical beauty is perhaps the most extreme aesthetic experience that is dependent upon culture and learning; those not versed in the language of mathematics cannot experience the beauty of a mathematical formulation. And yet, once the language of mathematics is mastered, the same formulae can be experienced as beautiful by mathematicians belonging to different races and cultures. Indeed, Paul Dirac coined the term "the principle of mathematical beauty” and made the beauty of a mathematical formula, rather than its simplicity, the ultimate guide to its truthfulness (Dirac 1939). He was not alone; other mathematicians, such as Hermann Weyl and Paul Erdös, thought similarly.

In what does the beauty of a mathematical formula lie? Perhaps the most forceful answers, and the ones nearest to our belief, come from Immanuel Kant on the one hand and Bertrand Russell on the other. Kant’s views are opaque and difficult to understand, and his use of the term "intuition” especially vague. For an interpretation of what constitutes mathematical beauty for Kant we rely on Breitenbach’s (2015) discussion. For Kant, it seems, a mathematical formulation is beautiful if it “makes sense”. This, of course, raises the question of “makes sense” to what. We believe that at least part of the experienced beauty of a mathematical formulation lies in the fact that it adheres to the logical deductive system of the brain and hence makes sense in terms of that logical deductive system. This is clearly stated by Russell (1920) in his *Introduction to Mathematical Philosophy*. Although he makes no reference to the brain, Russell equates mathematics with the brain’s logical deductive system when he asks *“What is this subject, which may be called indifferently either mathematics or logic?”* because, to him, *“What can be known, in mathematics and by mathematical methods, is what can be deduced from pure logic”* since *“logic is the youth of mathematics and mathematics is the manhood of logic”*. Perhaps most significantly for our argument, to Russell *“Logical propositions are such as can be known a priori, without study of the actual world”*. In other words, logical propositions can be traced to inherited brain concepts.

The logical deductive system of the brain, whatever its details, is inherited and is therefore similar in mathematicians belonging otherwise to different races and cultures. It is in this sense that mathematical beauty has its roots in a biologically inherited logical-deductive system that is similar for all brains. It is only by adhering to the rules of the brain’s logical deductive system that a formulation can gain universal assent and be found beautiful. Any departure from that would mean that it has lost the universal agreement. Implicit in our argument is that the experience of mathematical beauty, being the result of the application of the brain’s logical-deductive system, is a demonstration that the logical deductive system of mathematical brains, no matter what their cultural background may be, is the same. And since mathematical beauty, in our categorization, belongs to the biological category, it is not surprising that there is significantly less variability among mathematicians in rating mathematical equations as beautiful.

## Footnotes

3 When we sp, we restrict ourselves to saying that different humans do not differ when assigning different coloured patches, surfaces or objects to chips belonging to different colour categories. In saying so, we do not address the vexed question of colour qualia.

